# High-quality ultrastructural preservation using cryofixation for 3D electron microscopy of genetically labeled tissues

**DOI:** 10.1101/261594

**Authors:** Tin Ki Tsang, Eric A. Bushong, Daniela Boassa, Junru Hu, Benedetto Romoli, Sebastien Phan, Davide Dulcis, Chih-Ying Su, Mark H. Ellisman

**Author notes:** These authors contributed equally to this work (Tin Ki Tsang, Eric A. Bushong). Correspondence (M.H.E); (C.Y.S).

## Abstract

Electron microscopy (EM) offers unparalleled power to study cell substructures at the nanoscale. Cryofixation by high-pressure freezing offers optimal morphological preservation, as it captures cellular structures instantaneously in their near-native states. However, the applicability of cryofixation is limited by its incompatibilities with diaminobenzidine labeling using genetic EM tags and the high-contrast *en bloc* staining required for serial block-face scanning electron microscopy (SBEM). In addition, it is challenging to perform correlated light and electron microscopy (CLEM) with cryofixed samples. Consequently, these powerful methods cannot be applied to address questions requiring optimal morphological preservation and high temporal resolution. Here we developed an approach that overcomes these limitations; it enables genetically labeled, cryofixed samples to be characterized with SBEM and 3D CLEM. Our approach is broadly applicable, as demonstrated in cultured cells, *Drosophila* olfactory organ and mouse brain. This optimization exploits the potential of cryofixation, allowing quality ultrastructural preservation for diverse EM applications.

## INTRODUCTION

The answers to many questions in biology lie in the ability to examine the relevant biological structures accurately at high resolution. Electron microscopy (EM) offers the unparalleled power to study cellular morphology and structure at nanoscale resolution (Leapman, 2004). Cryofixation by high-pressure freezing (hereafter referred to as cryofixation) is the optimal fixation method for samples of thicknesses up to approximately 500 µm (Dahl and Staehelin, 1989; McDonald, 1999; Moor, 1987; Shimoni et al., 1998). By rapidly freezing the samples in liquid nitrogen (-196 °C) under high pressure (∼2100 bar), cryofixation immobilizes cellular structures within milliseconds and preserves them in their near-native states. In contrast, cross-linking based chemical fixation takes place at higher temperatures (≥4 °C) and depends on the infiltration of aldehyde fixatives, a process which takes seconds to minutes to complete. During chemical fixation, cellular structures may deteriorate or undergo rearrangement (Korogod et al., 2015; Steinbrecht and Müller, 1987; Szczesny et al., 1996) and enzymatic reactions can proceed (Kellenberger et al., 1992; Sabatini, 1963), potentially resulting in significant morphological artifacts.

Cryofixation is especially critical, and often necessary, for properly fixing tissues with cell walls or cuticles that are impermeable to chemical fixatives, such as samples from yeast, plant, *C. elegans*, and *Drosophila* (Ding, 1993; Doroquez et al., 2014; Kaeser et al., 1989; Kiss et al., 1990; McDonald, 2007; Müller-Reichert et al., 2003; Shanbhag et al., 1999, 2000; Winey et al., 1995). As cryofixation instantaneously halts all cellular processes, it also provides the temporal control needed to capture fleeting biological events in a dynamic process (Hess et al., 2000; Watanabe et al., 2013; Watanabe et al., 2013; Watanabe et al., 2014).

Despite the clear benefits of cryofixation, it is incompatible with diaminobenzidine (DAB) labeling reactions by genetic EM tags. For example, APEX2 (enhanced <u>a</u>scorbate <u>pe</u>ro<u>x</u>idase) is an engineered peroxidase that catalyzes DAB reaction to render target structures electron dense (Lam et al., 2015; Martell et al., 2012). Despite the successful applications of APEX2 to three-dimensional (3D) EM (Joesch et al., 2016), there has been no demonstration that APEX2 or other genetic EM tags can be activated following cryofixation. Conventionally, cryofixation is followed by freeze-substitution (Steinbrecht and Müller, 1987), during which water in the sample is replaced by organic solvents. However, the resulting dehydrated environment is incompatible with the aqueous enzymatic reactions required for DAB labeling by genetic EM tags.

EM structures can also be genetically labeled with fluorescent markers through correlated light and electron microscopy (CLEM). Yet, performing CLEM with cryofixed samples also presents challenges. Fluorescence microscopy commonly takes place either before cryofixation (Brown et al., 2009; Kolotuev et al., 2010; McDonald, 2009) or after the sample is embedded (Kukulski et al., 2011; Nixon et al., 2009; Schwarz and Humbel, 2009). However, if the specimen is dissected from live animals, the time taken to acquire fluorescence images delays cryofixation and could cause ultrastructural deterioration. In order for fluorescence microscopy to take place after embedding, special acrylic resins need to be used (Kukulski et al., 2011; Nixon et al., 2009; Schwarz and Humbel, 2009) and only a low concentration of osmium tetroxide stain can be tolerated (De Boer et al., 2015; Watanabe et al., 2011). These constraints limit the applicability of CLEM for cryofixed samples.

Another disadvantage of cryofixation is that *en bloc* staining during freeze-substitution is often inadequate. As a result, post-staining of ultramicrotomy sections is frequently needed for cryofixed samples (Shanbhag et al., 1999, 2000; Takemura et al., 2013). However, post-staining could be labor-intensive and time-consuming, especially for volume EM (Ryan et al., 2016; Zheng et al., 2017). Critically, on-section staining is impossible for samples imaged with block-face volume EM techniques (Briggman and Bock, 2012), such as serial block-face scanning electron microscopy (SBEM) (Denk and Horstmann, 2004). A large amount of heavy metal staining is necessary for SBEM to generate sufficient back-scatter electron signal and prevent specimen charging (Deerinck et al., 2010; Kelley et al., 1973; Tapia et al., 2012). Therefore, it remains impossible to image cryofixed samples with SBEM or other techniques that require high-contrast staining.

To overcome these limitations of cryofixation, here we present a robust approach, named the CryoChem Method (CCM), which combines key advantages of cryofixation and chemical fixation. This technique enables labeling of target structures by genetically encoded EM tags or fluorescent markers in cryofixed samples, and permits high-contrast *en bloc* heavy metal staining sufficient for SBEM. Specifically, we rehydrate cryofixed samples after freeze-substitution to make the specimen suitable for subsequent aqueous reactions and fluorescence imaging. We further show that 3D CLEM can be achieved by combining SBEM with confocal microscopy imaging of the frozen-rehydrated specimen. We successfully apply CCM to multiple biologically significant systems with distinct ultrastructural morphologies, including cultured mammalian cells, *Drosophila* olfactory organ (antenna) and mouse brain. By overcoming critical technical barriers, our method exploits the potential of cryofixation, making it compatible with genetically encoded EM tags, fluorescence imaging before resin embedding, and any EM techniques that require substantial heavy metal staining.

## RESULTS

Given that a key limitation of cryofixation arises from the dehydrated state of the samples after freeze-substitution (Figure 1), it is imperative that our approach delivers a cryofixed specimen that is fully hydrated and can then be processed at higher temperatures (4°C or room temperature) for enzymatic reactions and/or high-contrast *en bloc* heavy metal staining. It has been demonstrated that cryofixed samples can be rehydrated for immunogold labeling following cryosectioning (van Donselaar et al., 2007), but the method only yields modest EM contrast and is not compatible with genetic labeling using APEX2 nor volume EM techniques.

**Figure 1.**
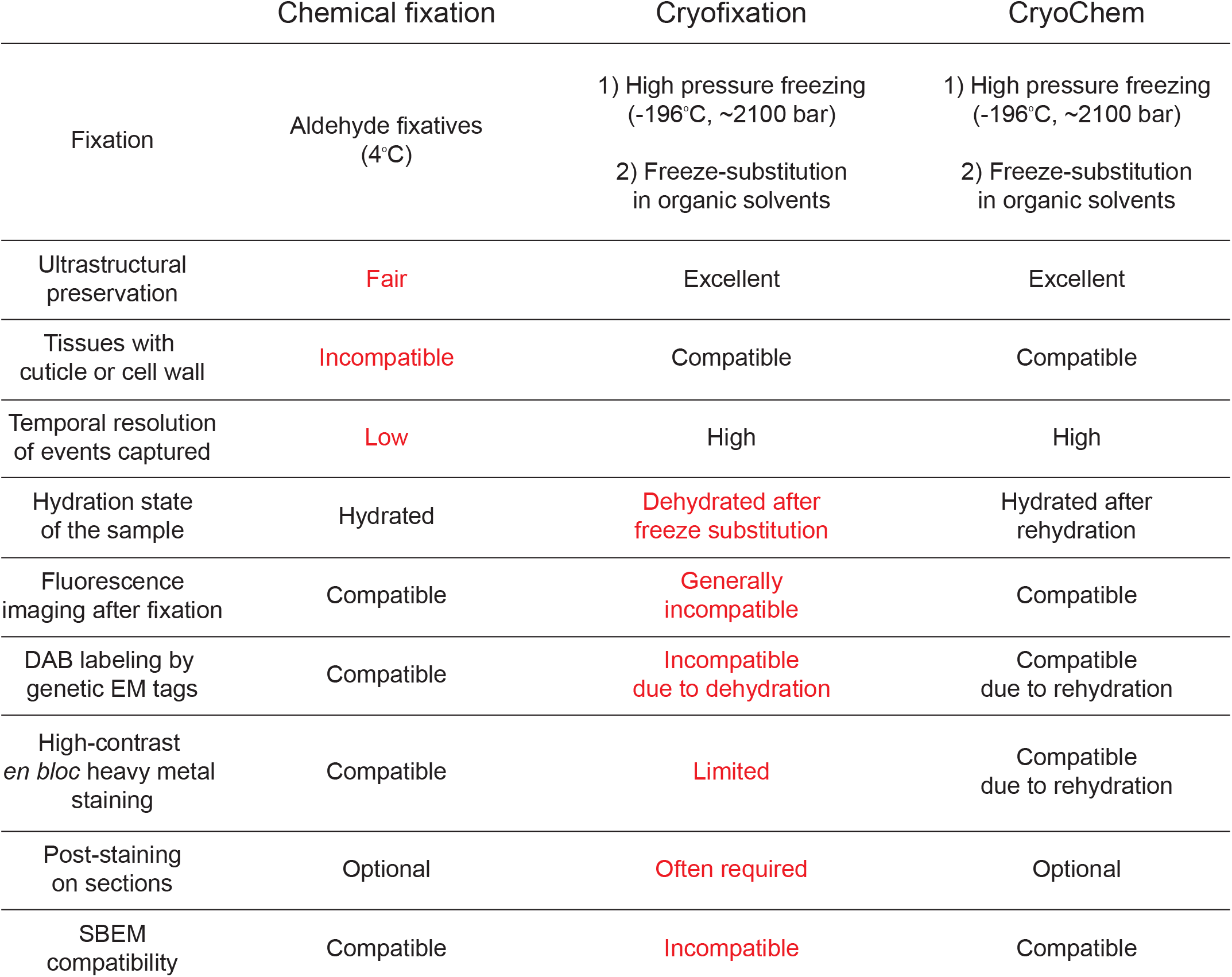
Comparison of the advantages and limitations of different sample preparation methods for electron microscopy. The CryoChem Method (CCM) combines the advantages of chemical fixation and cryofixation. With CCM, samples are fixed with high-pressure freezing and freeze-substitution to achieve quality ultrastructural preservation. This approach allows preservation of tissues with cuticle or cell wall and captures biological events with high temporal resolution. A rehydration step is introduced to enable fluorescence imaging, DAB labeling by genetically encoded EM tags and high-contrast *en bloc* heavy metal staining of the cryofixed sample. The high-contrast *en bloc* heavy metal staining permitted by CCM reduces the need for post-staining on sections, and makes CCM compatible with serial block-face scanning electron microscopy (SBEM). Common limitations of chemical fixation and cryofixation are denoted in red.

### The CryoChem Method

To achieve the ultrastructural preservation of cryofixation and the versatility of chemical fixation, we developed a hybrid protocol which we refer to hereafter as the CryoChem Method (CCM) (Figure 1). Importantly, we devised a freeze-substitution cocktail (see below) that allows preservation of APEX2 enzymatic activity and signals from fluorescent proteins. CCM begins with high-pressure freezing of the sample, followed by freeze-substitution in an acetone solution with glutaraldehyde (0.2%), uranyl acetate (0.1%), and water (1%), to further stabilize the cryo-preserved structures at low temperatures. After freeze-substitution, the sample is rehydrated gradually on ice with a series of acetone solutions containing an increasing amount of water or 0.1M HEPES. Once completely rehydrated, the cryofixed sample is amenable for imaging with fluorescence microscopy, DAB labeling reactions using genetically encoded tags, and the high-contrast *en bloc* staining (e.g. osmium-thiocarbohydrazide-osmium and uranyl acetate) normally reserved only for chemically fixed samples. Afterwards, samples are dehydrated through a series of ethanol solutions and acetone, then infiltrated with epoxy resin and cured using standard EM procedures. These resin embedded samples may be sectioned or imaged directly with any desired EM technique (Figure 2, see *Materials and Methods* for details).

**Figure 2.**
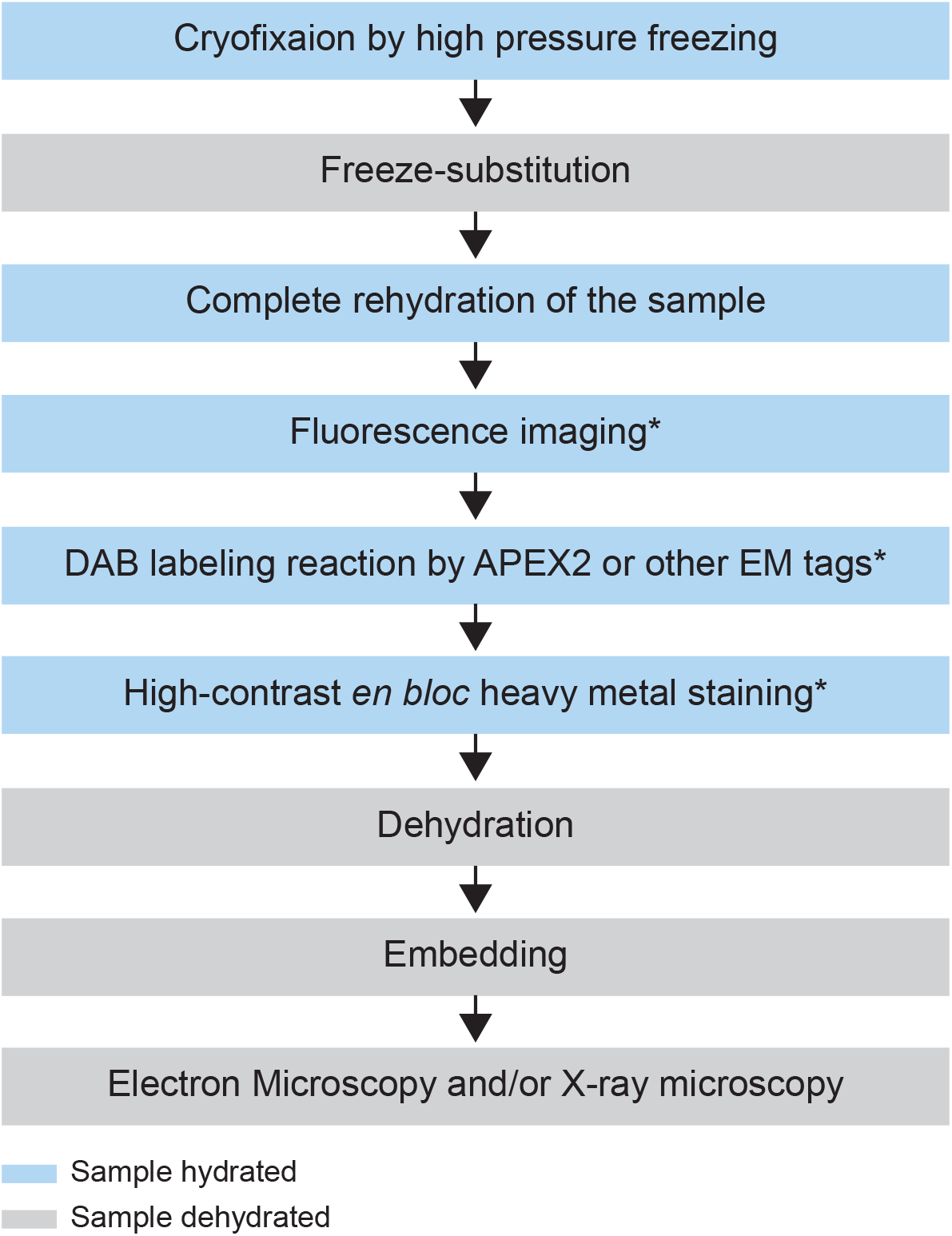
Flowchart of the CryoChem Method. After cryofixation by high-pressure freezing and freeze-substitution, cryofixed samples are rehydrated gradually. Rehydrated samples can then be imaged for fluorescence, subjected to DAB labeling reaction or *en bloc* stained with a substantial amount of heavy metals. The protocol is modular; the first three processes are the core steps of CCM and the starred steps are optional depending on the experimental design. The samples are then dehydrated for resin infiltration and embedding, followed by imaging with any EM technique of choice. Blue and grey denote hydrated and dehydrated states of the sample, respectively.

### CryoChem Method offers high-quality ultrastructural preservation and sufficient *en bloc* staining for SBEM

To determine whether CCM provides high-quality ultrastructural preservation, we first tested the method in a mammalian cell line. Using transmission electron microscopy (TEM), well-preserved mitochondria and nuclear membranes were observed in the CCM-processed cells (Figure 3— figure supplement 1). Given that cryofixation is often necessary for properly fixing tissues surrounded by a barrier to chemical fixatives, we next tested CCM in a *Drosophila* olfactory organ, the antenna, which is encased in a waxy cuticle (Figure 3A). A hallmark of optimally preserved antennal tissues prepared by cryofixation is the smooth appearance of membrane structures (Shanbhag et al., 1999, 2000; Steinbrecht, 1980; Steinbrecht and Müller, 1987). In the insect antenna, auxiliary cells extend microlamellae to surround the olfactory receptor neurons (ORNs), forming the most membrane-rich regions in the antenna. We therefore focused on this structure to evaluate the quality of morphological preservation afforded by our method. In CCM-processed antennal tissues, we found that the delicate structures of the microlamellae were well-preserved (Figure 3A and Figure 3—video supplement 1), unlike the chemically fixed counterparts in which microlamellae were distorted (Figure 3A) (Steinbrecht, 1980). Importantly, the overall ultrastructural preservation achieved through CCM resembles that obtained by standard cryofixation and freeze-substitution protocols (Shanbhag et al., 1999, 2000).

**Figure 3.**
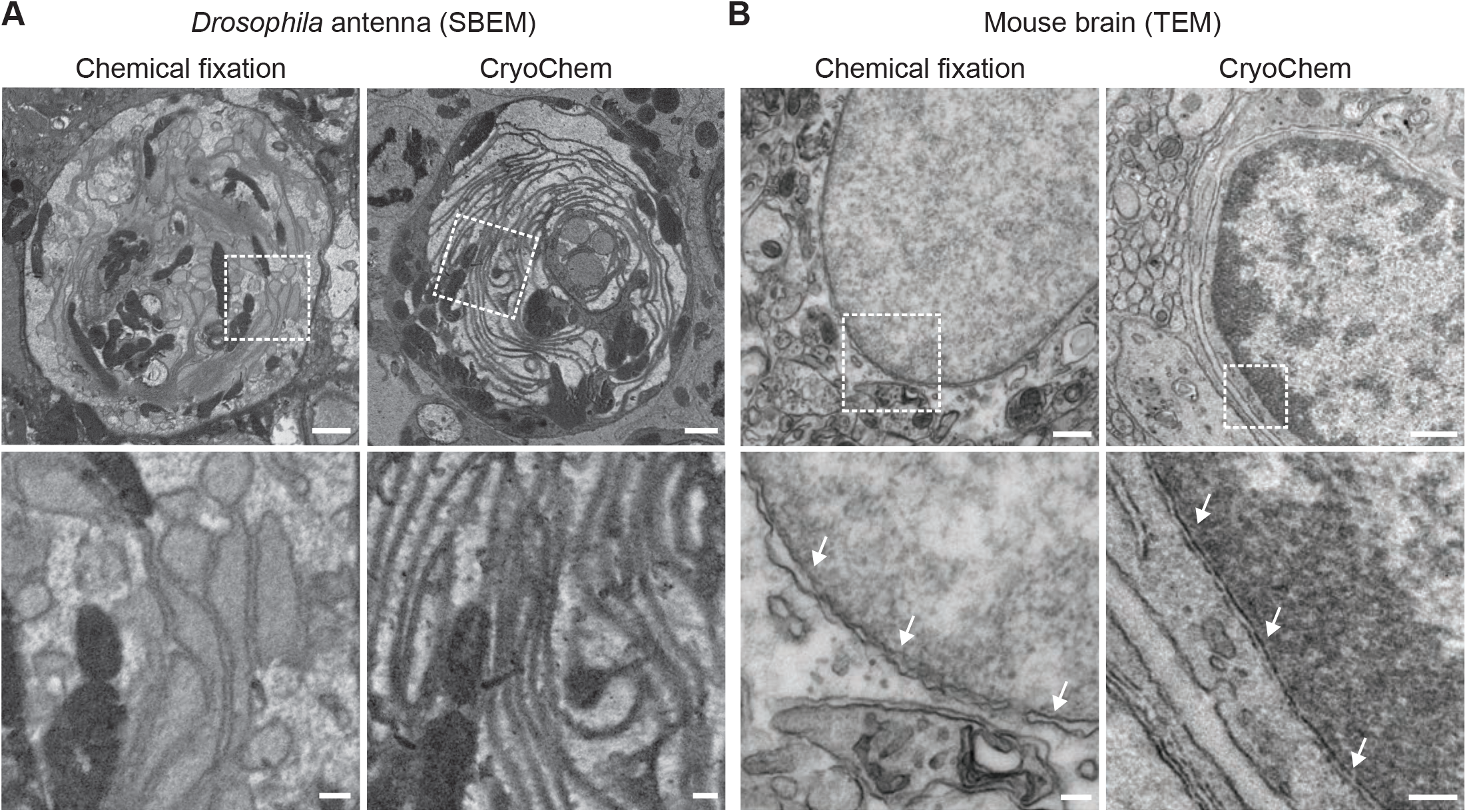
CryoChem Method offers high-quality ultrastructural preservation and sufficient *en bloc* staining for SBEM. TEM and SBEM images were acquired to assess the morphology of CCM-processed tissues. (A) The microlamella structures were well-preserved in CCM-processed *Drosophila* antenna (top right panel), compared to chemically fixed samples (top left panel). In the enlarged views of the boxed regions (bottom panels), the microlamellae in the CCM-processed antenna appeared uniform in size and shape, unlike the chemically fixed ones which were distorted. Scale bars: 1 μm for top panels, 200 nm for bottom panels. (B) CCM enhanced the morphological preservation of aldehyde-perfused mouse brain (top right panel), compared to the chemically fixed control (top left panel). In the enlarged views of the boxed regions (bottom panels), the nuclear membranes (arrows) are smoother in the CCM-processed sample. We note that the chromatin was more heavily stained in the CCM-processed specimen, likely due to the additional exposure to uranyl acetate during freeze substitution. Scale bars: 500 nm for top panels, 100 nm for bottom panels.

In contrast to fly antennae, which can be dissected expeditiously and frozen in the live state, certain tissues (e.g., mouse brain) are difficult to cryofix from life without tissue damage caused by anoxia or mechanical stress associated with dissection. In these cases, cryofixation can be performed after aldehyde perfusion and still produce quality morphological preservation (Sosinsky et al., 2008). To test whether CCM can improve morphological preservation of aldehyde-perfused samples, we cryofixed vibratome sections (100 μm) from an aldehyde-perfused mouse brain and processed the sample with CCM. As a control, we used standard chemical fixation procedures to process the vibratome sections of the same brain (see *Materials and Methods* for details). Compared to chemically fixed controls, the membranes of the CCM-processed samples were markedly smoother, indicating an improvement in morphological preservation (Figure 3B). This result agrees with our previous observation that cellular morphology can be markedly improved even when cryofixation is performed after aldehyde perfusion (Sosinsky et al., 2008).

Of note, we adopted a high-contrast *en bloc* staining protocol (Deerinck et al., 2010; Tapia et al., 2012; West et al., 2010; Williams et al., 2011) when processing the *Drosophila* antennae and mouse brain. An adequate level of heavy metals was incorporated into these cryofixed samples to allow for successful imaging by SBEM (Figure 3A and 4), even without nitrogen gas injection for charge compensation (Deerinck et al., 2017) (Figure 3– video supplement 1 and Figure 4D). This *en bloc* staining protocol is normally reserved only for chemically fixed tissues, but is now made compatible with cryofixed samples by CCM.

**Figure 4.**
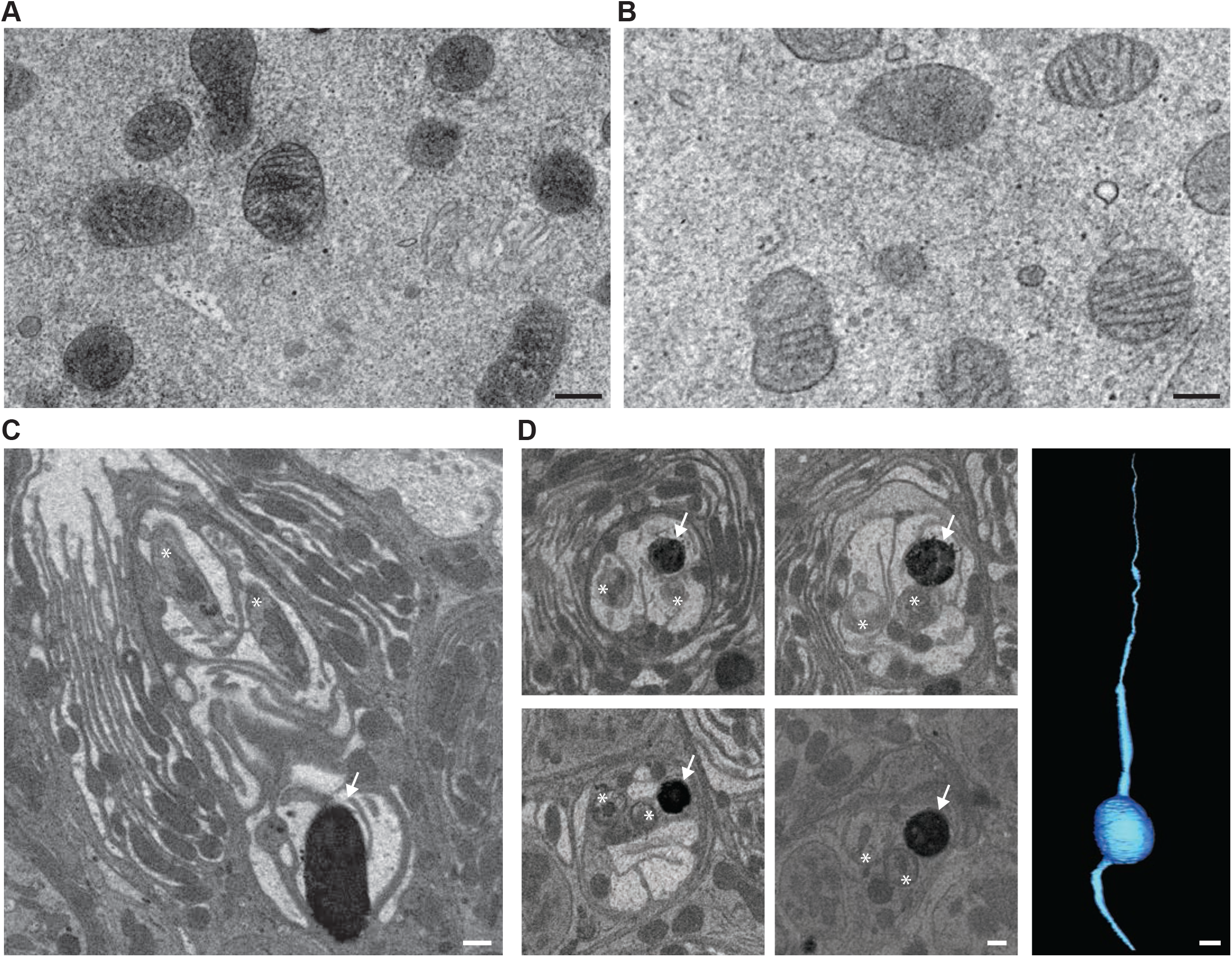
CryoChem Method enables DAB labeling by APEX2 in cryofixed tissues. In CCM-processed cultured cells and *Drosophila* antennae, DAB labeling was observed in cells expressing APEX2. (A) Mitochondria expressing APEX2 were labeled with DAB in a transfected HEK 293T cell. (B) An untransfected control cell. Scale bars: 200 nm. (C) An APEX2-expressing olfactory receptor neuron (ORN) was labeled with DAB (arrow) in the *Drosophila* antenna. Asterisks denote ORNs without APEX2 expression. Scale bar: 500 nm. (D) A series of SBEM images showing the same DAB labeled *Drosophila* ORN (arrow) in different planes of section. Asterisks denote ORNs without APEX2 expression. The images were acquired using standard imaging methods without charge compensation by nitrogen gas injection (Deerinck et al., 2017). These images, together with the rest of the EM volume acquired using SBEM, enabled semi-automatic segmentation and 3D reconstruction of the labeled ORN (right panel). Scale bars: 500 nm for SBEM images, 2 μm for the 3D model of ORN.

### CryoChem Method enables DAB labeling in cryofixed samples expressing APEX2

Next we determined if DAB labeling reaction can be performed in cryofixed samples with CCM. Using CCM-processed cultured cells expressing APEX2, we observed DAB labeling in the targeted organelles (mitochondria) in the transfected cells (Figure 4A), compared to the untransfected controls (Figure 4B). We further validated this approach in CCM-processed *Drosophila* antenna; successful DAB labeling was also detected in genetically identified ORNs expressing APEX2 with X-ray microscopy (Figure 4—video supplement 1). This imaging technique facilitates the identification of the region of interest for SBEM (Figure 4C), as we and others reported previously (Bushong et al., 2015; Ng et al., 2016). Crucially, we demonstrated that an EM volume of a genetically labeled, cryofixed ORN can be acquired with SBEM, which allows for an accurate 3D reconstruction of the ORN through semi-automated segmentation (Figure 4D). Taken together, these results demonstrate that CCM can reliably generate DAB labeling by genetically encoded EM tags in cryofixed samples.

### Fluorescence is well-preserved in CryoChem-processed samples

To determine whether CCM is compatible with fluorescence microscopy, we first evaluated the degree to which fluorescence level is affected after CCM processing. Using confocal microscopy, we quantified GFP fluorescence in the soma of unfixed *Drosophila* ORNs and that from CCM-processed samples after rehydration (Figure 5). Remarkably, GFP fluorescence intensities of fresh and CCM-processed ORNs are essentially indistinguishable with respect to their distributions (Figure 5A) and average levels (Figure 5B), indicating that CCM processing has little effect on GFP fluorescence in fly ORNs. Similarly, we observed strong GFP signals in the mouse brain after the cryofixed sample was rehydrated (Figure 5— figure supplement 1A).

**Figure 5.**
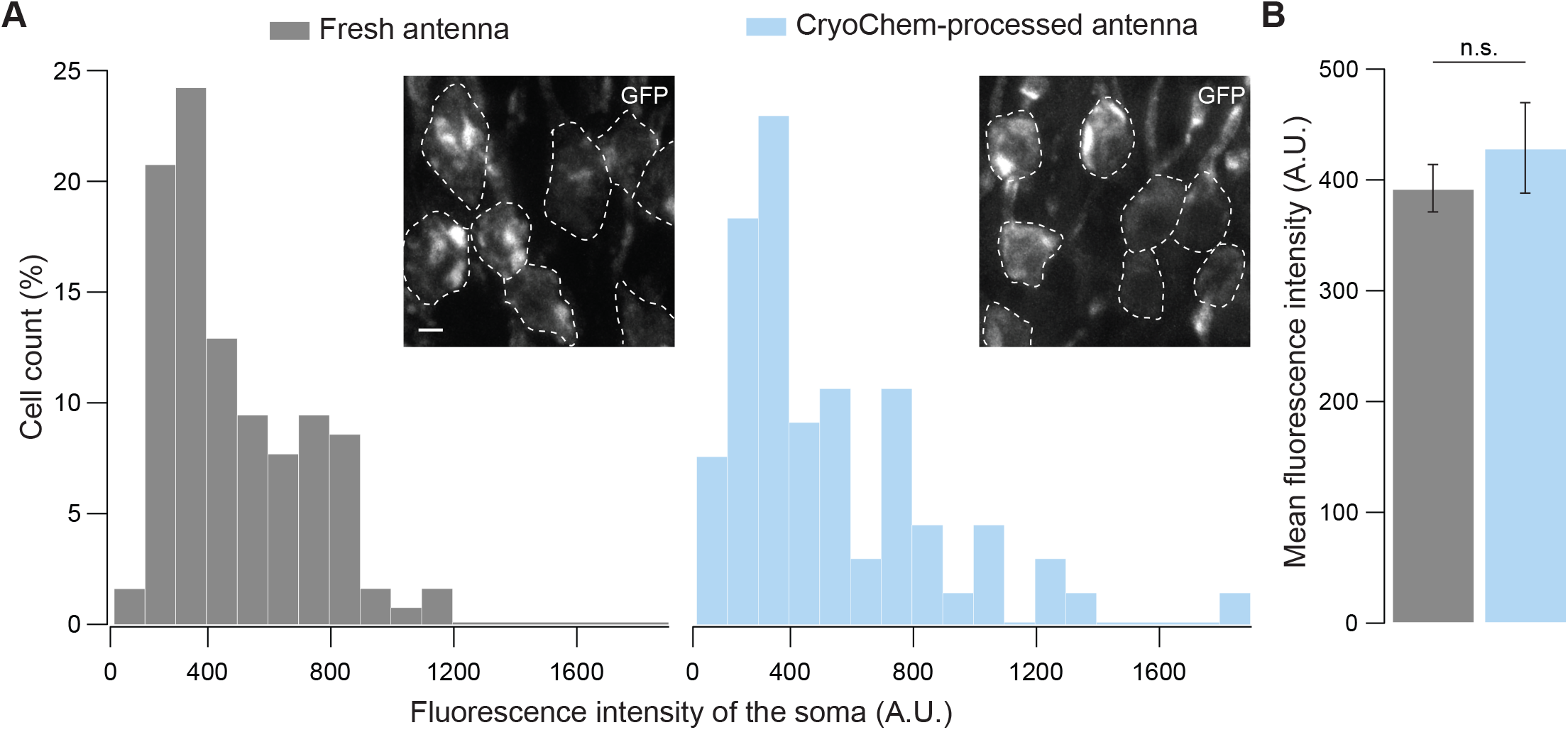
GFP fluorescence is well-preserved in CryoChem-processed samples. Confocal images were taken to quantify the level of GFP fluorescence in *Drosophila* ORNs. Antennae were collected from transgenic flies expressing GFP in a subset of ORNs. (A) GFP fluorescence intensity distributions of the ORN soma in freshly-dissected, unfixed antennae (left panel) and CCM-processed antennae (right panel) are not significantly different. *p* =0.810, Kolmogorov-Smirnov test. Insets show representative images, with ORN soma outlined. Scale bar: 2 μm. (B) Comparison of the average fluorescence intensities. GFP intensities are virtually identical between neurons in unfixed antennae and frozen-rehydrated antennae. n=3 antennae for each condition, Error bars denote SEM, *p* =0.950, Mann-Whitney *U* Test.

Next, we asked whether this observation also applies to another type of fluorescence protein. To this end, we examined tdTomato fluorescence in the mouse brain (Figure 5—figure supplement 1B). We note that tdTomato is not a variant of GFP and is instead derived from *Discosoma sp.* fluorescence protein ‘DsRed’ (Shaner et al., 2004). Confocal images of the CCM-processed mouse brain showed that the tdTomato fluorescence was also well-preserved (Figure 5—figure supplement 1B) and we were able to detect the co-expression of GFP and tdTomato in a subpopulation of neurons (Figure 5—figure supplement 1C). Taken together, our results indicate that CCM-processed sample can serve as a robust substrate for fluorescence imaging.

### 3D correlative light and electron microscopy (CLEM) in CCM-processed samples expressing fluorescent markers

Finally, we took advantage of the fact that fluorescence microscopy can now take place in a cryofixed sample before resin embedding to develop a protocol for 3D CLEM in CCM-processed specimens (Figure 6A, see *Materials and Methods* for details). The protocol first uses the core CCM steps to deliver a frozen-rehydrated sample. Subsequently, DRAQ5 DNA stain is introduced to the sample to label the nuclei, which can then serve as fiducial markers for CLEM. Next, the region containing target cells expressing fluorescent markers is imaged with confocal microscopy, during which signals from DRAQ5 and fluorescent markers are both acquired. After confocal microscopy, the sample is *en bloc* stained with multiple layers of heavy metals (Deerinck et al., 2010; Tapia et al., 2012; West et al., 2010; Williams et al., 2011), then dehydrated and embedded as in a typical CCM protocol. Subsequently, the embedded sample is imaged with X-ray microscopy. The resulting micro-computed tomography volume can be registered to the confocal volume using the nuclei as fiducial markers, so that the region of interest (ROI) for SBEM can be identified. After SBEM imaging, the EM volume can be registered to the confocal volume in a similar fashion for 3D CLEM.

**Figure 6.**
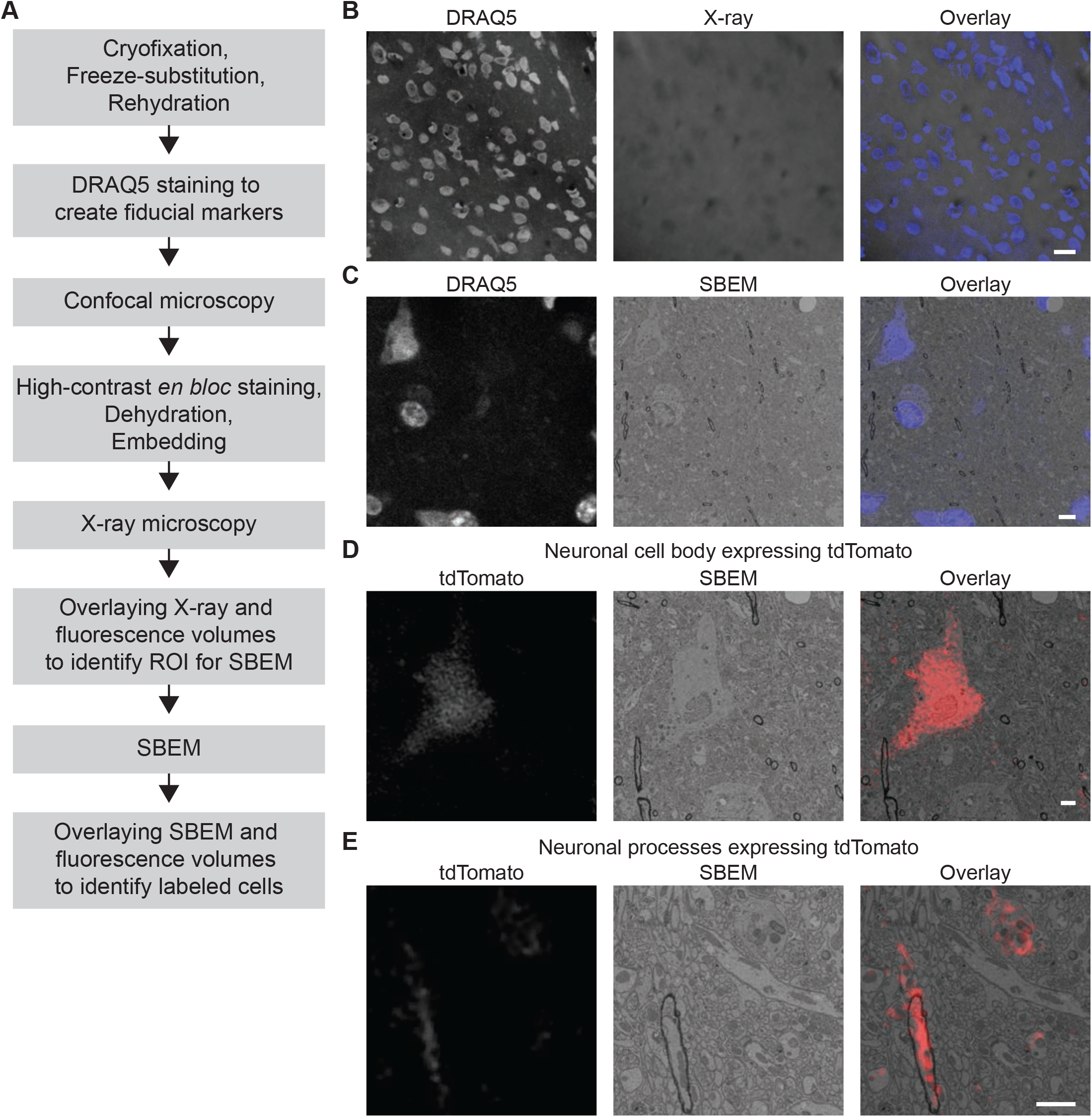
3D correlated light and electron microscopy (CLEM) in CCM-processed mouse brain. Mouse brain slices with fluorescently labeled neurons were processed with CCM, imaged with confocal microscopy, X-ray microscopy and SBEM for 3D CLEM. (A) Flowchart for performing 3D CLEM with CCM-processed samples. Similar to a typical CCM protocol, the cryofixed sample is first freeze-substituted and rehydrated. The frozen-rehydrated sample is then stained with DRAQ5 to label DNA in the nuclei. Next, the region of interest (ROI) is identified using confocal microscopy based on fluorescent signals, while the DRAQ5 signals are also acquired to serve as fiducial markers. Subsequently, the sample is stained, dehydrated and embedded for X-ray microscopy and SBEM. Using the DRAQ5 signals as fiducial markers, the confocal volumes can be registered to the X-ray volume such that the ROI for SBEM can be identified. Once the SBEM volume is acquired, it can be registered to the confocal volumes based on the positions of the nuclei for 3D CLEM. (B) An example of the DRAQ5 fluorescence signals (left), the corresponding ROI in X-ray volume (middle) and overlay (right). This image registration process facilitates ROI identification in SBEM. Scale bar: 20 μm. (C) DRAQ5 fluorescence labeling served as fiducial points for registering the confocal volume to the SBEM volume. Scale bar: 5 μm. (D) The cell body of a tdTomato-expressing neuron (left) was identified in the SBEM volume (middle) through 3D CLEM (right). (E) Neuronal processes expressing tdTomato (left) were also identified in the SBEM volume (middle) through 3D CLEM (right). Scale bars: 2 μm, for both (D) and (E).

As a proof of principle, we performed 3D CLEM in an aldehyde-perfused, CCM-processed mouse brain expressing tdTomato in a subset of neurons. To this end, we first determined if DRAQ5 staining can be performed in a frozen-rehydrated specimen. Using confocal microscopy, we were able to observe DRAQ5 labeling of the nuclei in a cryofixed brain slice after rehydration (Figure 6B). We used the labeled nuclei as fiducial markers to register the X-ray volume with the confocal data (Figure 6B) and thereby target a ROI with tdTomato-expressing neurons for SBEM imaging.

Similarly, we were able to register the confocal volume to the SBEM volume (Figure 6C). Of note, the CLEM accuracy was ensured by using the bright heterochromatin labeling by DRAQ5 and their corresponding structures in EM as finer fiducial points (Figure 6C). Furthermore, the fluorescent markers made it possible to identify the target cell bodies (Figure 6D and Figure 6 —video supplement 1) and small neuronal processes (Figure 6E and Figure 6—video supplement 1) in the SBEM volume. Lastly, we note that with CCM, fluorescence microscopy in cryofixed specimens can take place before *en bloc* EM staining. Therefore, our protocol does not require special resins for embedding and permits high-contrast staining with high concentrations of osmium tetroxide.

## DISCUSSION

We described here a hybrid method, named CryoChem, which combines key advantages of cryofixation and chemical fixation to substantially broaden the applicability of the optimal fixation technique. With CCM, it is now possible to label target structures with DAB by a genetically encoded EM tag, image cells expressing fluorescent markers before resin embedding and deposit high-contrast *en bloc* staining in cryofixed tissues. In addition, with CCM, one can also perform 3D CLEM in cryofixed specimens. Our method thereby provides an important alternative to conventional cryofixation and chemical fixation methods.

The modular nature of CCM (Figure 2) makes it highly versatile as researchers can modify the modules to best suit their needs. For instance, to prevent over-staining, one can replace the high-contrast *en bloc* staining step (osmium-thiocarbohydrazide-osmium and uranyl acetate) (Deerinck et al., 2010; Tapia et al., 2012; West et al., 2010; Williams et al., 2011) with a single round of osmium tetroxide staining for thin section TEM (Figures 3B, 4A and 4B) or electron tomography. In addition, CCM is essentially compatible with a wide range of reactions catalyzed by EM tags other than APEX2 (Ellisman et al., 2015). For example, the protein labeling reactions mediated by miniSOG (Shu et al., 2011) and the tetracysteine-based methods using FIAsH and ReAsH (Gaietta et al., 2002), or the non-protein biomolecule labeling reactions using Click-EM (Ngo et al., 2016) or ChromEM (Ou et al., 2017). The versatility of CCM will likely expand the breath of biological questions that can be addressed using cryofixed samples.

In addition to using EM tags, we have also developed a 3D CLEM protocol (Figure 6A) that allows optimally and rapidly preserved EM structures to be genetically labeled with fluorescent markers in CCM-processed tissues. In contrast to EM tags, fluorescent markers do not generate electron-dense products (e.g. DAB polymers) that can obscure the subcellular structures. Moreover, with multicolor CLEM, one can utilize multiple readily available genetically encoded fluorescent markers to label different target structures or cells. Using the 3D CLEM protocol, one could also pinpoint labeled subcellular structures (e.g., microtubules) or proteins (e.g., ion channels) in an EM volume with super-resolution microscopy.

The advantages of CCM makes it particularly suited for addressing biological questions that require optimal and rapid preservation of a genetically labeled structure. For example, to construct an accurate model to describe the biophysical properties of a neuron, it is essential to acquire morphological measurements based on faithfully preserved ultrastructures. CCM processing provides such an opportunity; we were able to obtain a 3D reconstruction of a genetically labeled *Drosophila* ORN at nanoscale resolution with quality morphological preservation (Figure 4D). In addition, by combining CCM with Flash-and-Freeze EM (Watanabe et al., 2014) and electron tomography, it is possible to capture the fast morphological changes of genetically labeled vesicles in 3D during synaptic transmission.

Furthermore, CCM is applicable to addressing questions in diverse tissue types, as demonstrated here with cultured mammalian cells or tissues of *Drosophila* antennae and mouse brains. Notably, identical solutions and experimental conditions were used for these different tissues in all core steps (Figure 2). Thus, the protocol described here can likely be readily adapted to cells and tissues of other biological systems. In addition, we demonstrated that CCM can further improve the ultrastructure of an aldehyde-perfused brain compared to chemically fixed counterparts (Figure 3B). Given that aldehyde perfusion is often required for the dissection of deeply embedded or fragile tissues, the compatibility of CCM with aldehyde fixation further broadens the applicability of the method.

## MATERIALS AND METHODS

### Cultured cells preparation

HEK 293T cells were grown on 1.2 mm diameter punches of Aclar (2 mil thick; Electron Microscopy Sciences, Hatfield, PA) for 48 hours, in a humidified cell culture incubator with 5% CO_2_ at 37 °C. The culture medium used was DMEM (Mediatech Inc., Manassas, VA) supplemented with 10% fetal bovine serum (Gemini Bio-Products, West Sacramento, CA). The cells were transfected with Lipofectamine 2000 (Invitrogen, Carlsbad, CA) with a plasmid carrying APEX2 targeted to mitochondria (pcDNA3-Mito-V5-APEX2, Addgene #72480) (Lam et al., 2015). At 24 hours after transfection, the cells were used for CCM processing.

### DNA constructs and *Drosophila* transgenesis

Orco cDNA was a gift from Dr. Aidan Kiely, and APEX2 cDNA was acquired from Addgene (APEX2-NES, #49386). Membrane targeting of APEX2 was achieved by fusing the marker protein to the C-terminus of mCD8GFP or to the N-terminus of Orco. Briefly, gel-purified PCR fragments of mCD8GFP, APEX2, and/or Orco were pieced together with Gibson Assembly following manufacturer’s instructions (New England Biolabs, Ipswich, MA). A linker (SGGGG) was added between APEX2 and its respective fusion partner. In the APEX2-Orco construct, a myc tag was included in the primer and added to the N-terminus of APEX2 to enable the detection of the fusion protein by immunostaining. To facilitate Gateway Cloning (ThermoFisher Scientific, Waltham, MA), the attB1 and attB2 sites were included in the primers and added to the ends of the Gibson assembly product by PCR amplification. The PCR products were then purified and cloned into pDONR221 vectors via BP Clonase II (Life Technologies, Carlsbad, CA). The entry clones were recombined into the pBID-UASC-G destination vector (Wang, Beck and McCabe, PloS ONE, 2012) using LR Clonases II (Life Technologies, Carlsbad, CA).

*Drosophila* transgenic lines were derived from germline transformations using the ϕC31 integration systems (Groth, Fish, Nusse, & Calos, 2004; Markstein, Pitsouli, Villalta, Celniker, & Perrimon, 2008). All transgenes described in this study were inserted into the attP40 landing site on the second chromosome (BestGene Inc., Chino Hills, CA). Target expression of APEX2 in the ORNs was driven by the Or47b-GAL4 driver (#9984, Bloomington Drosophila Stock Center, Figures 3-5) or the Or22a-GAL4 driver (Dobritsa et al. 2003, Figure 4—video supplement 1). Flies were raised on standard cornmeal food at 25 °C in a 12:12 light-dark cycle.

### *Drosophila* antennae preparation

Six to eight days old flies were cold anesthetized and then pinned to a Sylgard dish. The third segments of the antennae were removed from the head of the fly with a pair of fine forceps and then immediately transferred to a drop of 1X PBS on the dish. With a sharp glass microelectrode, a hole was poked in the antenna to facilitate solution exchange. It is critical that the tissue remained in PBS at all times to prevent deflation. The antenna should remain plump and maintain its shape prior to cryofixation.

### Chemical fixation of *Drosophila* antenna

Antennae were dissected as described above, and then incubated in Karnovsky fixatives (2% paraformaldehyde/2.5% glutaraldehyde/2 mM CaCl_2_ in 0.1 M sodium cacodylate) at 4°C for 18 hours. Next, samples were washed in 0.1 M sodium cacodylate for 10 minutes and in a solution of 100 mM glycine (Bio-Rad Laboratories, Hercules, CA) in 0.1 M sodium cacodylate for another 10 minutes, and twice more in 0.1 M sodium cacodylate. All washing steps were performed on ice. The following *en bloc* heavy metal staining, dehydration and resin embedding steps were carried out as described in the CryoChem Method section below.

### Transgenic mice and virus-mediated gene transfer

Animals were handled in accordance with the guidelines established by the *Guide for Care and Use of Laboratory Animals* of the National Institutes of Health and approved by the Animal Care and Use Committee of University of California, San Diego. To introduce GFP and tdTomato fluorescent markers in a mouse brain (Figure 5—figure supplement 1), GFP was expressed in the tyrosine hydroxylase (TH)-expressing neurons and tdTomato in the corticotropin releasing factor (CRF)-expressing neurons. A CRF driver mouse line (B6.Cg-Crh^tm1(cre)Zjh^/J, Jackson laboratory) expressing CRE recombinase under the control of the Crh promoter/enhancer elements was first crossed to a tdTomato reporter line (B6.Cg-Gt.ROSA.26Sor^tm14(CAG-tdTomato)Hze^/J, Jackson Laboratory). The progeny was then crossed to a TH-GFP mouse line (Kessler, Yang, Gollomp, Jin, & Iacovitti, 2003), obtaining a transgenic model stably expressing GFP in dopaminergic (TH^+^) neurons and CRE/tdTomato in CRF-releasing neurons. To test the 3D CLEM protocol and the morphological preservation offered by CCM (Figure 3B, Figure 6 and Figure 6—video supplement 1), a mouse brain from a tdTomato reporter line (B6.Cg-Gt.ROSA.26Sor^tm14(CAG-tdTomato)Hze^/J, Jackson Laboratory) was used.

### Mouse brain preparation

Mice were anesthetized with ketamine/xylazine and then transcardially perfused with Ringer’s solution followed by 0.15 M sodium cacodylate containing 4% paraformaldehyde/0.2% (Figure 5—figure supplement 1) or 0.5% (Figure 3B, Figure 6 and Figure 6—video supplement 1) glutaraldehyde/2 mM CaCl_2_. The animal was perfused for 10 minutes with the fixatives and then the brain was removed and placed in ice-cold fixative for 1 hour. The brain was then cut into 100-μm thick slices using a vibrating microtome. Slices were either processed for chemical fixation (Figure 3B) or stored in ice-cold 0.15 M sodium cacodylate for around 4 hours until used for high-pressure freezing (Figure 3B, Figure 5—figure supplement 1, Figure 6 and Figure 6— video supplement 1).

### Chemical fixation of mouse brain

The aldehyde-perfused mouse brain slices were post-fixed in 2.5% glutaraldehyde for 20 minutes, then washed with 0.15 M sodium cacodylate five times for 5 minutes on ice. Next, the samples were incubated in 0.15 M sodium cacodylate with 100 mM glycine for 5 minutes on ice, then washed in 0.15 M sodium cacodylate similarly. The following *en bloc* heavy metal staining, dehydration and resin embedding steps were carried out as described in the CryoChem Method section below.

#### CryoChem Method

##### (I) Cryofixation by high-pressure freezing

###### Cultured cells

Aclar disks were placed within the well of a 100 µm-deep membrane carrier. The cells were covered with the culture medium and then high pressure frozen with a Leica EM Pact 2 unit.

###### Drosophila antennae

The third antennal segment was dissected as described above. Antennae from the same fly were transferred into the 100 µm-deep well of a type A planchette filled with 20% BSA (Sigma-Aldrich, St. Louis, MO) in 0.15 M sodium cacodylate. The well of the type A planchette was then covered with the flat side of a type B planchette to secure the sample. The samples were immediately loaded into a freezing holder and frozen with a high-pressure freezing machine (Bal-Tec HPM 010). Planchettes used for cryofixation were pre-coated with 1-hexadecene (Sigma-Aldrich, St. Louis, MO) to prevent planchettes A and B from adhering to each other so as to allow solution to reach the samples during freeze-substitution.

###### Mouse brain slices

A 1.2 mm tissue puncher was used to cut a portion of hypothalamus expressing tdTomato (Figure 3B, Figure 5—figure supplement, Figure 6 and Figure 6—video supplement 1) and GFP (Figure 5—figure supplement) from a tissue slice. The tissue punch was placed into a 100 μm-deep membrane carrier and surrounded with 20% BSA in 0.15 M sodium cacodylate. The specimen was high-pressure frozen as described for the *Drosophila* antennae.

All frozen samples were stored in liquid nitrogen until further processing.

##### (II) Freeze-substitution

Frozen samples in a planchette were transferred to cryo-vials containing the following freeze-substitution solution: 0.2% glutaraldehyde (#18426, Ted Pella, Redding, CA), 0.1% uranyl acetate (Electron Microscopy Sciences, Hatfield, PA), and 1% water in acetone (#AC326800010, ACROS Organics, USA) in a liquid nitrogen bath. The sample vials were then transferred to a freeze-substitution device (Leica EM AFS2) at −90 °C for 58 hours, from −90 °C to −60 °C for 15 hours (with the temperature raised at 2 °C/hr), at −60 °C for 15 hours, from −60 °C to −30 °C for 15 hours (at +2 °C/hr), and then at −30 °C for 15 hours. In the last hour at −30°C, samples were washed three times in an acetone solution with 0.2% glutaraldehyde and 1% water for 20 minutes. The cryo-tubes containing the last wash were then transferred on ice for an hour.

##### (III) Rehydration

The freeze-substituted samples were then rehydrated gradually in a series of nine rehydration solutions (see below). The samples were transferred from the freeze-substitution solution to the first rehydration solution (5% water, 0.2% glutaraldehyde in acetone) on ice for 10 minutes. The rehydration step was repeated in a stepwise manner until the samples were fully rehydrated in the final rehydration solution (0.1 M and 0.15 M sodium cacodylate for cells and antennae or mouse brain slices, respectively) (van Donselaar et al., 2007):

1. 5% water, 0.2% glutaraldehyde in acetone
2. 10% water, 0.2% glutaraldehyde in acetone
3. 20% water, 0.2% glutaraldehyde in acetone
4. 30% water, 0.2% glutaraldehyde in acetone
5. 50% 0.1M HEPES (Gibco, Taiwan), 0.2% glutaraldehyde in acetone
6. 70%, 0.1M HEPES, 0.2% glutaraldehyde in acetone
7. 0.1 M HEPES
8. 0.1 M / 0.15 M sodium cacodylate with 100 mM glycine
9. 0.1 M / 0.15 M sodium cacodylate

After rehydration, samples were removed from the planchettes using a pair of forceps under a stereo microscope to a 0.1 M / 0.15 M sodium cacodylate solution in a scintillation vial on ice. It is important that subsequent DAB labeling and *en bloc* heavy metal staining are carried out in scintillation vials instead of the planchettes because metal planchettes may react with the labeling or staining reagents.

##### (IV) DRAQ5 staining

Mouse brain slices were incubated in DRAQ5 (1:1000 in 0.15 M sodium cacodylate buffer; ThermoFisher Scientific, Waltham, MA) on ice for 60 minutes. Then the samples were washed in 0.15 M sodium cacodylate three times for 10 minutes on ice before fluorescence imaging.

##### (V) Fluorescence imaging

###### Drosophila antenna

Freshly dissected or cryofixed-rehydrated antennae (*10x UAS-APEX2-mCD8GFP; Or47b-GAL4*) were mounted in FocusClear (elExplorer, Taiwan) between two cover glasses (#1.5 thickness, 22 mm x 22 mm, Fisher Scientific, Hampton, NH) separated by two layers of spacer rings. Confocal images were collected on an Olympus FluoView 1000 confocal microscope with a 60X water-immersion objective lens. The 488 nm laser was used to excite GFP and all images were acquired at the same laser power and gain to enable comparison between the fresh vs cryofixed-rehydrated samples.

###### Mouse brain slices

Confocal images of mouse brain slices were collected on a Leica SPE II confocal microscope with a 20X water-immersion objective lens using 488 nm and 561 nm excitation. After freeze-substitution and rehydration, the specimens were placed in ice-cold 0.15 M sodium cacodylate for imaging. Confocal volumes of DRAQ5 and tdTomato signals were collected on an Olympus FluoView 1000 confocal microscope with 20X air and 60X water objectives.

##### (VI) DAB labeling of target structures by APEX2

###### Cultured cells

Samples were transferred to a 0.05% DAB (#D5637, Sigma-Aldrich, St. Louis, MO) solution in 0.1 M sodium cacodylate for 5 minutes on ice to allow DAB to diffuse into the tissue. To label the mitochondria in the APEX2-expressing cells, samples were then transferred to a 0.05% DAB solution with 0.015% H_2_O_2_ (Fisher Scientific, Hampton, NH) in 0.1 M sodium cacodylate until DAB labeling was visible under a microscope (∼5 minutes on ice). After the reaction, samples were washed three times with 0.1 M sodium cacodylate on ice for 10 minutes.

###### Drosophila antennae

Samples were first placed into a 0.05% DAB solution in 0.1 M sodium cacodylate for an hour on ice to allow DAB to access target neurons underneath the cuticle in the antenna. To label APEX2-expressing ORNs, antennae were then transferred into a 0.05% DAB solution with 0.015% H_2_O2 in 0.1 M sodium cacodylate for an hour on ice. After the reaction, samples were washed three times with 0.1 M sodium cacodylate on ice for 10 minutes.

##### (VII)*En bloc* heavy metal staining for TEM and SBEM

For TEM: Cultured cells and mouse brain slices were incubated in 2% OsO_4_ (Electron Microscopy Sciences, Hatfield, PA) /1.5% potassium ferrocyanide (Mallinckrodt, Staines-Upon-Thames, UK) /2 mM CaCl_2_ in 0.1 M (cells) or 0.15 M (brain) sodium cacodylate for an hour on ice. Then samples were washed in water five times for 5 minutes on ice prior to the dehydration step detailed below.

For SBEM: *Drosophila* antennae and mouse brain slices were incubated in 2% OsO_4_/1.5% potassium ferrocyanide/2 mM CaCl_2_ in 0.1 M (antennae) or 0.15 M (brain) sodium cacodylate for an hour at room temperature. Then samples were washed in water five times for 5 minutes and transferred to 0.5% thiocarbohydrazide (filtered with 0.22 μm filter before use; Electron Microscopy Sciences, Hatfield, PA) for 30 minutes at room temperature. Samples were washed in water similarly and incubated in 2% OsO_4_ for 30 minutes at room temperature. Afterwards, samples were rinsed with water, then transferred to 2% aqueous uranyl acetate (filtered with 0.22 μm filter) at 4 °C overnight. In the next morning, samples were first washed in water five times for 5 minutes and then subjected to the dehydration steps detailed below.

##### (VIII) Dehydration

Samples were dehydrated with a series of ethanol solutions and acetone in six steps of 10 minutes each: 70% ethanol, 90% ethanol, 100% ethanol, 100% ethanol, 100% acetone, 100% acetone. All ethanol dehydration steps were carried out on ice, and the acetone steps at room temperature. The first acetone dehydration step was carried out with ice-cold acetone, and the second one was with acetone kept at room temperature.

##### (IX) Resin infiltration

###### Cultured cells

Samples were transferred to a Durcupan ACM resin/acetone (1:1) solution for an hour on a shaker at room temperature. The samples were then transferred to fresh 100% Durcupan ACM resin overnight and subsequently placed in fresh resin for four hours. While in 100% resin, samples were placed in a vacuum chamber on a rocker to facilitate the removal of residual acetone. Finally, the samples were embedded in fresh resin at 60 °C for two days.

###### Drosophila antennae and mouse brain slices

Samples were transferred to a Durcupan ACM resin/acetone (1:1) solution overnight on a shaker. The next day, samples were transferred into fresh 100% Durcupan ACM resin twice, with six to seven hours apart. While in 100% resin, samples were placed in a vacuum chamber on a rocker to facilitate the removal of residual acetone. After the overnight incubation in 100% resin, samples were embedded in fresh resin at 60 °C for at least two days.

Durcupan ACM resin (Sigma Aldrich, St. Louis, MO) composition was 11.4 g component A, 10 g component B, 0.3 g component C, and 0.1 g component D.

### X-ray Microscopy (microcomputed tomography)

#### Drosophila antenna

Microcomputed tomography (microCT) was performed on resin-embedded specimens using a Versa 510 X-ray microscope (Zeiss). Flat-embedded specimens were glued to the end of an aluminum rod using cyanoacrylic glue. Imaging was performed with a 40X objective using a tube current of 40 kV and no source filter. Raw data consisted of 1601 projection images collected as the specimen was rotated 360 degrees. The voxel dimension of the final tomographic reconstruction was 0.4123 μm.

#### Mouse brain slices

X-ray microscopy scan was collected of a resin-embedded sample at 80 kVp with a voxel size of 0.664 μm prior to mounting for SBEM imaging. A second scan was collected of the mounted specimen at 80 kVp with 0.7894 μm voxels.

### Transmission Electron Microscopy

Ultrathin sections (70 nm) were collected on 300 mesh copper grids. Samples were post-stained with either Sato’s lead solution only (cultured cells) or with 2% uranyl acetate and Sato’s lead solution (mouse brain slices). Sections were imaged on an FEI Spirit TEM at 80 kV equipped with a 2k x 2k Tietz CCD camera.

### Serial Block-face Scanning Electron Microscopy

#### Drosophila antenna

Following microcomputed tomography to confirm proper orientation of region of interest, specimens were mounted on aluminum pins with conductive silver epoxy (Ted Pella, Redding, CA). The specimens were trimmed to remove excess resin above ROI and to remove silver epoxy from sides of specimen. The specimens were sputter coated with gold-palladium and then imaged using a Gemini scanning electron microscope (Zeiss) equipped with a 3View2XP and OnPoint backscatter detector (Gatan). Images were acquired at 2.5 kV accelerating voltage with a 30 μm condenser aperture and 1 μsec dwell time; Z step size was 50 nm. Volumes were either collected in variable pressure mode with a chamber pressure of 30 Pa and a pixel size of 3.8 nm (Figure 3—video supplement 1 and Figure 4D) or using local gas injection (Deerinck et al., 2017) set to 85% and a pixel size of 6.5 nm (Figures 3A and 4C). Volumes were aligned using cross correlation, segmented, and visualized using IMOD.

#### Mouse brain slices

SBEM was performed on a Merlin scanning electron microscope (Zeiss) equipped with a 3View2XP and OnPoint backscatter detector (Gatan). The volume was collected at 2 kV, with 6.8 nm pixels and 70 nm Z steps. Local gas injection (Deerinck et al., 2017) was set to 15% during imaging. The raster size was 10k x 15k and the Z dimension was 659 sections.

### Semi-automated segmentation of DAB-labeled *Drosophila* olfactory receptor neuron

The DAB-labeled *Drosophila* ORN was segmented in a semi-automated fashion using the IMOD software (Kremer, Mastronarde, & McIntosh, 1996) to generate the 3D model. The IMOD command line ‘imodauto’ was used for the auto-segmentation by setting thresholds to isolate the labeled cellular structures of interest. Further information about the utilities of ‘imodauto’ can be found in the IMOD manual (http://bio3d.colorado.edu/imod/doc/man/imodauto.html). Auto-segmentation was followed by manual proofreading and reconstruction by two independent proofreaders. The proofreaders used elementary operations in IMOD, most commonly the ‘drawing tools’ to correct the contours generated by ‘imodauto’. Where ‘imodauto’ failed to be applied successfully, the proofreaders also used the ‘drawing tools’ to directly trace the outline of the labeled structure. The contours of ORNs generally do not vary markedly between adjacent sections. Therefore, alternate sections were traced for the reconstruction of some parts of the ORN dendrite.

### Quantification of fluorescence intensity

To quantify GFP fluorescence intensity shown in Figure 5, maximum intensity Z-projections were generated using ImageJ (NIH). Average fluorescence intensity in the background was subtracted from the fluorescence intensity of each cell body measured. Only non-overlapping cell bodies were quantified. Kolmogorov-Smirnov Test was performed on http://www.physics.csbsju.edu/stats/KS-test.html and Mann-Whitney *U* Test was performed using SigmaPlot 13.0 (Systat Software, San Jose, CA).

### Light and electron microscope volume registration

To target tdTomato-expressing cells in the mouse brain for SBEM imaging, the confocal volumes collected in the frozen-rehydrated specimen was registered with the microCT volume of the resin-embedded sample, using a software tool developed in our lab. The resin-embedded specimen was then mounted and trimmed for SBEM based on the microCT volume. A second microCT scan of the mounted specimen allowed for precise targeting of the cells of interest with the Gatan stage for SBEM. After the SBEM volume was collected, the confocal and SBEM volumes were registered using the landmark tool of Amira 6.3 (ThermoFisher, Waltham, MA). Heterochromatin structures revealed by DRAQ5 labeling and visible in the SBEM volume were used as landmark points for the registration.

## ACKNOWLEDGEMENTS

We thank Aiden Keily for providing the Orco cDNA construct and R. Alexander Steinbrecht for advice on electron microscopy of *Drosophila* antennae. We also thank Edie Zhang and Martin Orden for assistance in segmentation of *Drosophila* olfactory receptor neuron. We also thank Andrea Thor and Mason Mackey for help with EM sample preparation and imaging. We also thank Steven Wasserman for comments on the manuscript.

---

**Figure supplement 1 for Figure 3.**
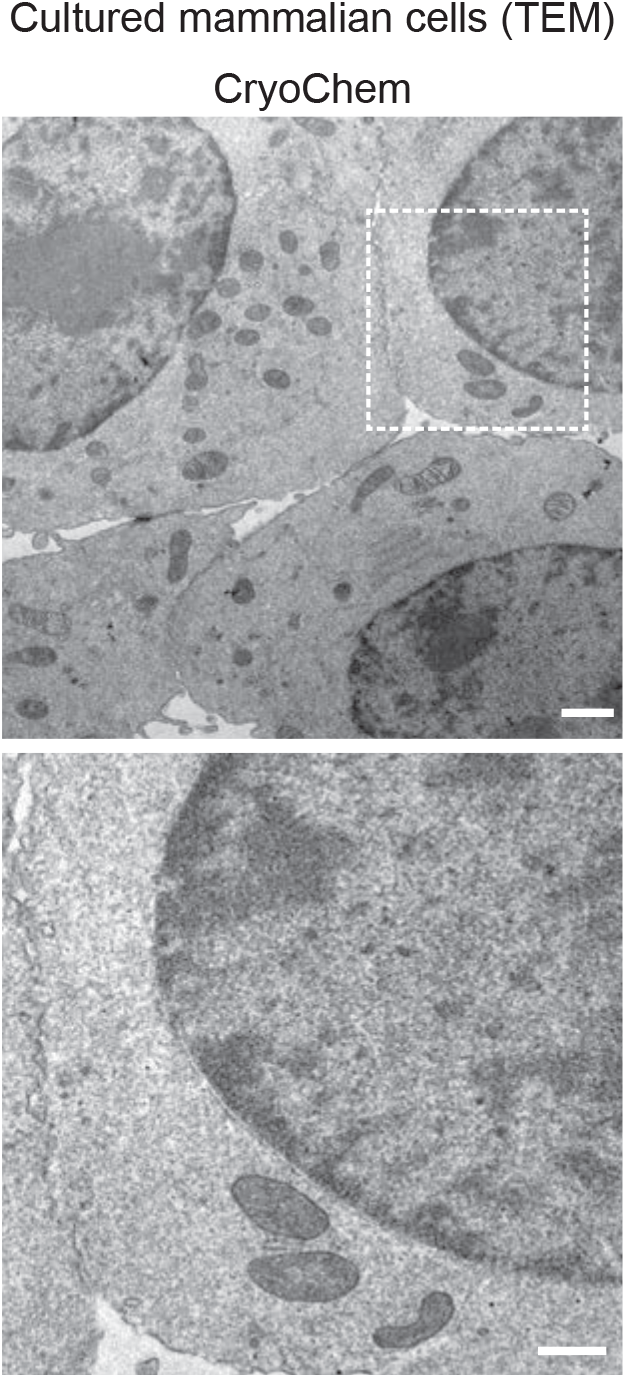
TEM images showed well-preserved ultrastructures in the CCM-processed HEK 293T cells. In the enlarged view of the boxed region (bottom panel), smooth nuclear membranes were observed. Scale bars: 1 μm for the top panel, 500 nm for the bottom panel.

**Figure supplement 1 for Figure 5.**
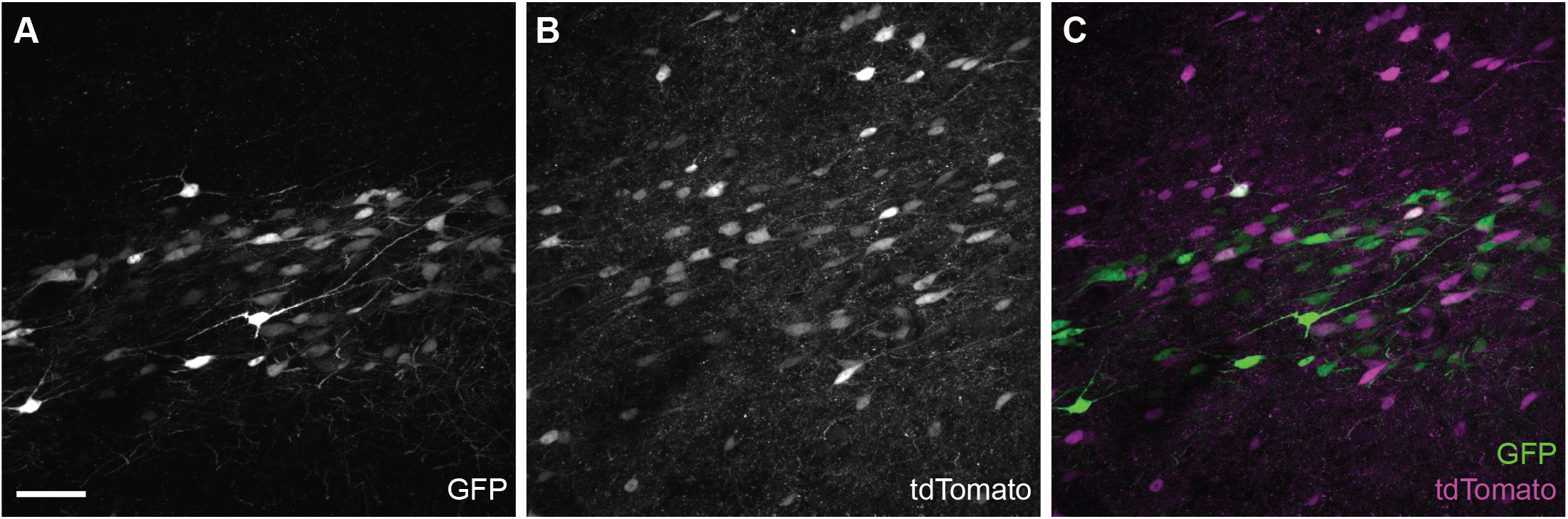
GFP and tdTomato fluorescence in a cryofixed-rehydrated mouse brain. (A) GFP- and (B) tdTomato-positive neurons. (C) Co-expression of GFP and tdTomato fluorescence was detected in the cryofixed-rehydrated mouse brain. Scale bar: 50 μm.

**Video supplement 1 for Figure 3**. A SBEM volume from a CryoChem-processed *Drosophila* antenna. Scale bar: 500 nm.

**Video supplement 1 for Figure 4**. An X-ray micro-computed tomography volume from a CCM-processed *Drosophila* antenna showing DAB labeling in subsets of ORNs expressing APEX2. The damaged region on the opposite side of the labeled cells indicates the hole poked in the antenna to facilitate solution exchange. Scale bar: 30 μm.

**Video supplement 1 for Figure 6**. A volume showing 3D CLEM in a CCM-processed mouse brain. tdTomato-expressing neurons were clearly identified in the SBEM volume. Scale bar: 10 μm.

## SOURCE DATA FILES

**Figure 5-Source Data 1** GFP fluorescence intensities in fresh and CCM-processed *Drosophila* antennae.

